# IKK2/NFkB signaling controls lung resident CD8 T cell memory during influenza infection

**DOI:** 10.1101/2022.02.15.480615

**Authors:** Curtis J. Pritzl, Dezzarae Luera, Karin M. Knudson, Michael J. Quaney, Mark A. Daniels, Emma Teixeiro

## Abstract

CD8 tissue resident memory (T_RM_) cells are especially suited to control pathogen spread at mucosal sites. However, their maintenance in lung is limited. Here, we found that enhancing NFkB signaling in T cells once memory to influenza is established increased pro-survival Bcl-2 and CD122 levels boosting lung CD8 T_RM_ maintenance. By contrast, enhancing NFkB signals during the contraction phase of the response led to a defect in T_RM_ differentiation without impairing recirculating memory subsets. Specifically, inducible activation of NFkB via constitutive active IKK2 or tumor necrosis factor (TNF) interfered with tumor growth factor beta (TGFβ) signaling resulting in defects of lung CD8 T_RM_ imprinting molecules CD69, CD103, Runx3 and Eomes. Conversely, inhibiting NFkB signals not only recovered but improved the transcriptional signature and generation of lung CD8 T_RM_. Thus, NFkB signaling is a critical regulator of tissue resident memory, whose levels can be tuned at specific times during infection to boost lung CD8 T_RM_.

## Introduction

Once infection has resolved, a few of the pathogen specific T cells that participated in the response persist as memory cells providing the host with enhanced protection against re-infection (1-3). These memory T cells strategically relocate to blood and secondary lymphoid organs (central, T_CM_ and effector, T_EM_, memory) as well as portal of entry tissues (tissue resident, T_RM_) each, with specific phenotypes and functions (4). Together, they guarantee the generation of a diverse and polyfunctional T cell memory pool. In contrast to other memory subsets, T_RM_ cells do not leave tissue, and continue patrolling it for signs of pathogen re-entry. If this happens, they trigger innate immune responses and immediately control reinfection in situ, in tissues like lung, skin or gut(5). T_RM_ cells have a protective role not only in infectious diseases(6-9), but also in cancer (10-13). Yet, mounting evidence also associates T_RM_ with pathology in autoimmunity, transplants, and graft versus host disease (14-16). Although this puts T_RM_ as a therapeutical target to treat disease, there is still poor understanding of how T_RM_ cells are generated or maintained in tissues. Furthermore, the times during the immune response that are suitable for manipulation of T_RM_ for therapeutic purposes are still ill defined. This is particularly important in the case of respiratory infections such as influenza that depend on lung-CD8 T_RM_ to control viral titers and disease severity(17, 18) but where CD8 T_RM_ longevity is limited(18).

One of the cardinal features of T_RM_ cells is their imprinting of non-lymphoid “tissue residency”, which differentiates them from circulating T cells. This is phenotypically characterized by high expression of CD69 and often (but not always) CD103. Transcriptionally, CD8 T_RM_ cells require high expression of Runx3(11), Nr4a1 (19, 20) and low expression of Eomes (21), although depending on the tissue, a balanced expression of other transcription factors, such as Blimp-1 in lung(22), is also important. Signals that occur prior to tissue entry (23) and tissue-specific signals (24) both contribute to the differentiation of T_RM_. Among these, antigen and tumor growth factor β (TGFβ) signals act at different points of the immune response to shape T_RM_(25-31). Yet, the role of inflammation in the generation and maintenance of T_RM_ remains largely unexplored.

NFkB signaling is a major driver of inflammation (32, 33) as well as one of the signaling pathways induced by T cell receptor signaling upon antigen recognition(34). Multiple pro-inflammatory factors (such as TNF or IL-1 or TLRs), together with antigen, signal through the canonical NFkB pathway at different times during infection (35-37), making it a plausible signaling hub where different environmental cues converge to regulate T cell differentiation and cell fate decisions. Here we sought to understand how changes in the levels of IKK2/NFkB signaling a CD8 T cell experiences during infection impact their memory fate. Our data show that NFkB signaling has a specific role in tissue resident memory that is different from the other recirculating memory subsets. Furthermore, NFkB signaling differentially regulates CD8 T_RM_ differentiation and CD8 T_RM_ maintenance. Interestingly, our data also reveals that tuning NFkB signaling levels at specific times during influenza infection can aid to boost or deplete CD8 T_RM_ in the lung, an organ where these cells gradually vanished over time after vaccination or infection leading to loss in clinical protection(18, 38).

## Results

### Increasing the levels of NFkB signaling after the peak of the response improves circulating CD8 T cell memory

To address the impact of NFkB signaling on T cell protective immunity, we generated two tetON inducible systems restricted to the T cell lineage. For this, we crossed mice carrying either a constitutively active Ikbkb allele (CA-IKK2)(39) or a dominant negative-acting version of IKK2 (DN-IKK2)(40) driven by the tetracycline TA-activated promoter (tetO)7 transactivator with mice expressing CD2-driven rtTA(41). We refer to these mice as CD2rtTAxCA-IKK2 and CD2rtTAxDN-IKK2 respectively (*SI Appendix* Fig. S1 and S2). Expression of CA- and DN-IKK2 can be monitored by a luciferase reporter (either by flow or by luciferase assays) and is restricted to the T cell lineage (*SI Appendix* Fig.S1 and S2C). Furthermore, doxycycline dependent induction of IKK2 results in the upregulation of NFkB dependent genes (CD69 and Eomes)(42, 43) but does not lead to overt T cell apoptosis (no induction of cleaved caspase-3 or FasL) (*SI Appendix* Fig. S2D).

We used these two new inducible models to interrogate whether changing the levels of IKK2/NFkB signaling in T cells during specific phases of the immune response impacts CD8 T cell memory. We tested whether boosting (or inhibiting) IKK2/NFkB signal transduction could modulate the establishment of circulating CD8 memory in two different polyclonal models of infection. For this, we used both tetON IKK2 inducible models and manipulated NFkB signaling after the peak of the T cell response following on previous reports suggesting a role for p65NFkB transcriptional activity during the contraction phase of the immune response to *L. monocytogenes* (42, 44). We found that inhibition of NFkB signaling, during the contraction phase of the immune response to influenza infection, led to a loss of circulating CD8 T cell memory. By contrast, increasing NFkB signaling resulted in a considerable increase in the number of polyclonal antigen specific memory CD8 T cells generated, both against influenza A (IAV) and VSV virus infection (*SI Appendix* Fig.S2E-H). From these data, we concluded that IKK2/NFkB signaling has a critical role in the establishment of CD8 T cell memory upon infection and most importantly, that increasing the levels of this signaling pathway in T cells at the end of the immune response improves the generation of CD8 T cell memory cells.

### IKK2/NFkB signaling differentially regulates T cell memory subset diversity

Since CD8 T cell memory is composed of different subsets with unique locations and phenotypes, we next asked whether the impact of IKK2/NFkB signaling would equally affect all T cell memory subsets. Using our inducible models, we infected mice with X31 IAV and allowed CD8 T cells to differentiate for 5 days after which, we either increased or decreased NFkB signaling in T cells by feeding mice with doxycycline chow during the contraction phase of the immune response. Then, we measured the frequency and number of IAV specific CD8 T_CM_, T_EM_ and T_RM_ generated (Fig. 1A). Inhibition of NFkB signaling during contraction, impaired the generation of both T_CM_ and T_EM_. However, increasing NFkB signaling, improved only the generation of IAV specific T_CM_ (Fig. 1B, C). Most strikingly, when we examined the generation of lung T_RM_, we observed that NFkB signaling had opposite effects on this T cell subset. Generation of NP_366_ and PA_224_ IAV antigen specific lung CD8 T_RM_ was decreased under high levels of NFkB signals. On the contrary, decreasing NFkB signaling resulted in a dramatic boost in the generation of polyclonal lung IAV (NP and PA) specific CD8 T_RM_ (Fig 1D,E). This was not exclusive to the lung or the type of infection as the same results were observed for IAV specific CD8 T_RM_ in spleen (Fig. 1F) and kidney in the context of systemic VSV infection (Fig. 1G). Of note, increasing NFkB levels did not result in an overall decrease in the number of CD8 or CD4 T cells in the lung, indicating that the effects of increasing NFkB signaling levels were restricted to antigen specific CD8 T cells (Fig. 2B,C). We also did not observe important defects in the lung IAV-specific CD4 T cell memory compartment when NFkB signals were increased, suggesting that the impairment in CD8 T_RM_ generation we found was not due to a defect in CD4 T cells (Fig. 2D). Finally, we used an adoptive transfer model to confirm whether the effect of NFkB signaling was CD8 T cell intrinsic. In response to both, influenza and i.n. VSV infection we found that increasing NFkB signaling specifically in CD8 T cells during the contraction phase led to a severe loss in antigen specific lung CD8 T_RM_ (Fig. 2E,F). Altogether these data reveal that IKK2/NFkB signaling is a critical pathway in the regulation of CD8 T cell memory subset diversity. Most importantly, our data suggest that manipulating NFkB signaling could deplete or boost CD8 T_RM_ in tissue.

**Figure 1.**
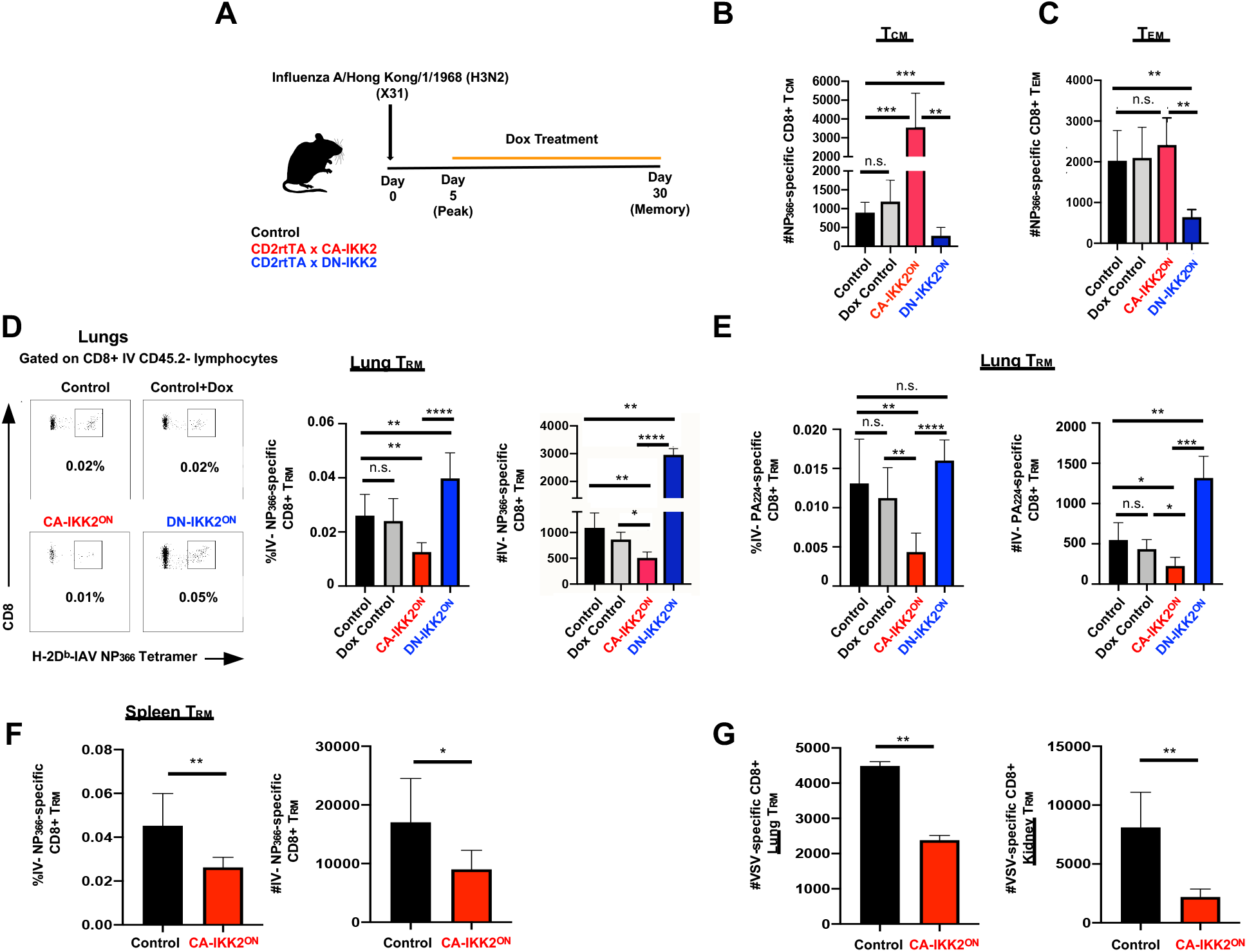
NFkB signaling differentially regulates T cell memory subset diversity. Groups of control, CD2rtTA x CA-IKK2 or CD2rtTA x DN-IKK2 mice (n≥3 mice per group) were infected with influenza x31. From day 5-30 p.i., mice were fed a doxycycline containing diet (control littermates+dox, CA-IKK2^ON^ or DN-IKK2^ON^ CD8 T cells) or control diet (CD2rtTA x CA-IKK2; CD2rtTA x DN-IKK2 or control littermates fed with regular chow) (A). At 30 days p.i., the numbers of influenza NP_366-374_-specific central (CD8+ Db-NP-tet+ CD44^hi^ CD62L^hi^ T_CM_) (B) and effector (CD8+ Db-NP-tet+ CD44^hi^ CD62L^lo^ T_EM_) (C) memory subsets were distinguished by flow cytometry in mediastinal lymph nodes. (D-F) Resident memory T cells (T_RM_) were identified in x31-infected mice by intravascular staining with CD45.2-specific, PE-labeled antibody injection prior to euthanasia. Frequencies and numbers of NP_366-374_-specific (D) and PA_224-233_-specific (E) CD8 T_RM_-cells were determined in lungs (Db-NP/PA-tetramers+ CD8+ CD45.2-) by flow cytometry. (F) Frequencies and numbers of NP_366-374_-specific CD8+ T_RM_ in the spleens. Representative data shown from 3 independent experiments. (G) Groups of control or CD2rtTA x CA-IKK2 (CA-IKK2^ON^) mice were infected with VSV. Mice were fed a doxycycline-containing diet from days 5 – 30 p.i.. At 30 days p.i., VSV-specific CD8+ T_RM_ (Kb-N-tet+ CD8+ CD45.2-) were identified in the lungs and kidneys of infected mice by iv. labelling. Representative data from 3 independent experiments. * p < 0.05, ** p < 0.01, *** p < 0.001, n.s. not significant.

**Figure 2.**
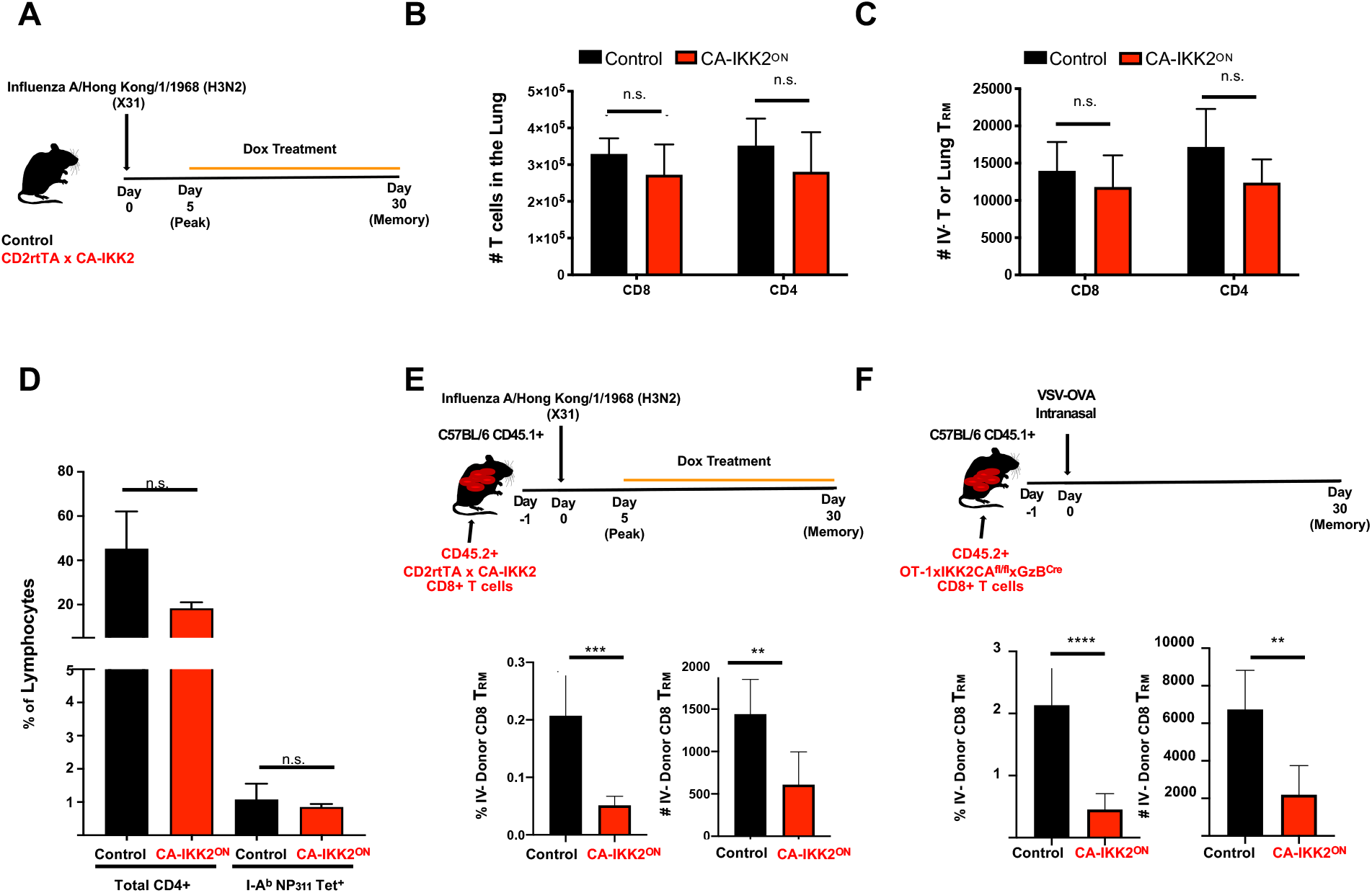
Impairment in the generation of CD8 T_RM_ under enhanced NFkB signals is T cell intrinsic. (A) Groups of control or CD2rtTA x CA-IKK2 mice were infected intranasally with influenza x31 (15000 pfu). From day 5-30 p.i, mice were fed a doxycycline containing diet (CA-IKK2^ON^) or control diet. (B-C) Number of total and resident CD8 and CD4 memory T cells (T_RM_) in the lungs of infected mice, the latter determined by intravascular staining. (D) Total and Influenza virus NP_311-325_-specific CD4 positive T cells determined by flow cytometry upon Influenzax31 infection (5000pfu). (E) CD8^+^ naïve T cells from CD2rtTAxCA-IKK2 mice were adoptively transferred to congenic host mice. The recipients were infected with influenza x31. Groups of recipients were fed doxycycline-containing or control diets from days 5-30 p.i. At day 30 p.i., frequency and number of lung CD8^+^ donor T_RM_ cells were determined i by intravenous labeling with anti-CD8b PE-labelled antibodies. (F) OT-1 naïve CD8+ T cells isolated from OT-1xIKK2CA^fl/fl^xGzB^Cre^ mice were adoptively transferred to congenic hosts followed by intranasal VSV-OVA infection. At 30 days p.i., lung donor OT-1 cells were identified by intravascular staining. n ≥ 2 experiments with n≥3 mice per group. ** p < 0.01, *** p < 0.001, n.s. not significant.

### Increasing IKK2 signaling impairs protection against heterologous infections

CD8 T_RM_ are critical to provide protection in tissue against re-infection. We, thus, tested whether increasing the amount of NFkB signaling in CD8 T cells as they differentiate to T_RM_ would impact protective immunity in the lung. For this, we followed a published approach to deplete circulating CD8 T cells while sparing CD8 T_RM_. (45). We adoptively transferred OT-1 naïve male T cells into female or male congenically marked hosts, followed by intranasal VSV-OVA infection (Fig. 3A). Consistent with rejection against male antigen, male donors vanished in female but not in male hosts (Fig. 3B,C). These conditions allowed us to singularly evaluate the T cell protective ability of antigen specific lung CD8 T_RM_ generated under high levels of NFkB signaling. Consistent with our results in Fig. 2, high levels of NFKB signaling (CA-IKK2^ON^ model) led to a loss of CD8 T_RM_ (Fig. 3D). However, female hosts bearing only male CA-IKK2^ON^ CD8 T_RM_ cells exhibited ∼200 times higher virus titers than their control counterparts upon heterologous infection in the lung. This was despite better effector function (Fig 3E,F). These data, thus, show that loss of CD8 T_RM_ due to high levels of NFkB signaling impairs protective immunity in the lung.

**Figure 3.**
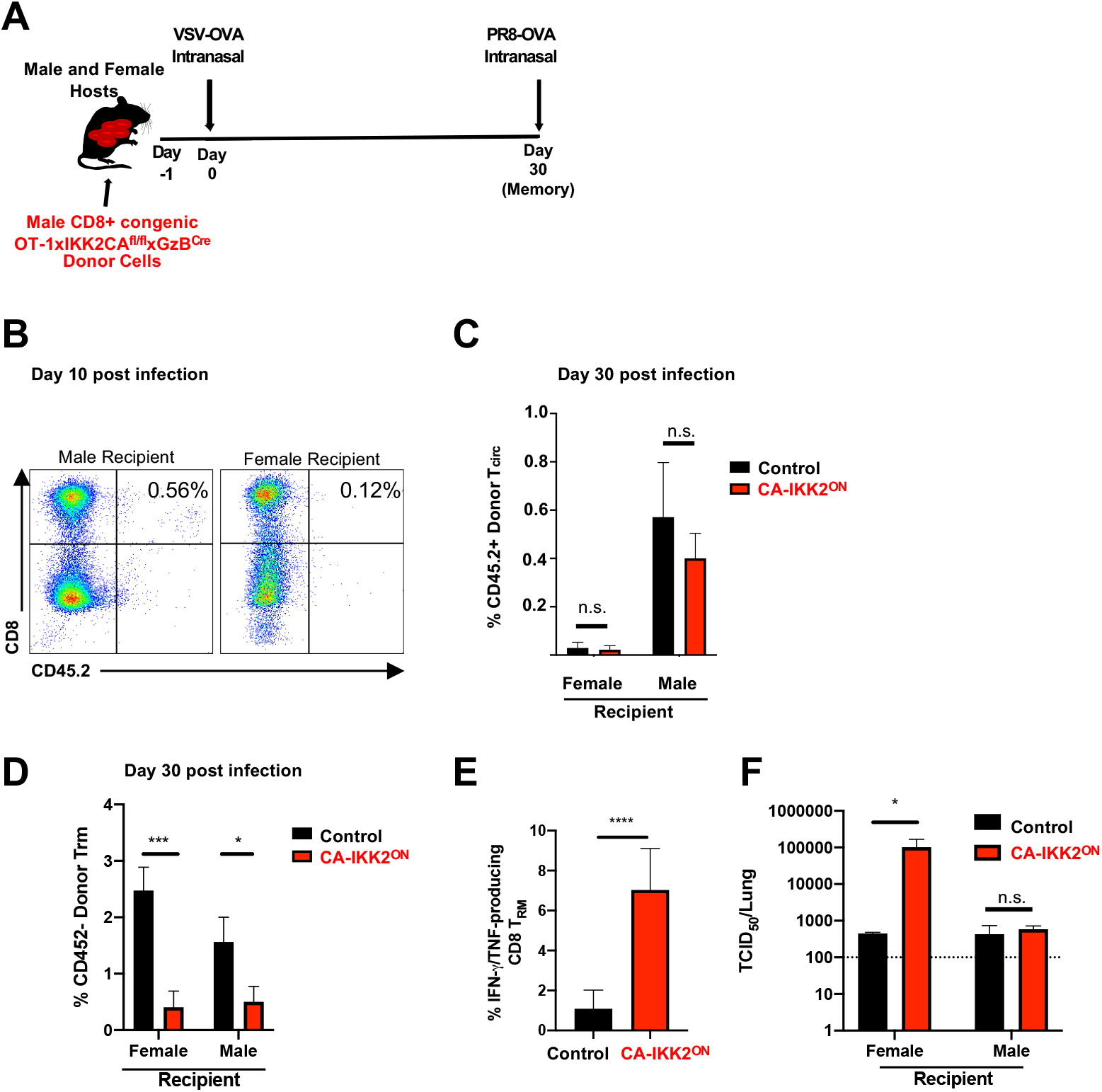
Enhancing NFkB signaling reduces T_RM_ and compromises protection against heterologous infection. (A) CD8 naive T cells from male OT-1xIKK2CA^fl/fl^xGzB^Cre^ or OT-1 littermate control naïve cells were adoptively transferred into groups of congenically marked male and female hosts, followed by intranasal VSV-OVA infection. Rejection of circulating cells was determined in all mice at day 10 p.i. (B). Frequency of circulating donor OT-1 CD8 T cells was assessed in the mediastinal lymph nodes at day 30 p.i., (C) and lung-resident, donor CD8 T-cells were determined by intravascular staining. (D). Frequency of IFNg and TNF double positive OT-1 CD8^+^ T_RM_ expressors at day 30 p.i. upon *ex vivo* antigen stimulation (E). A subset of mice from each group (n=5) were challenged with influenza PR8-OVA. Virus titers in the lungs were determined 2.5 days post-challenge (f). Data shown in pooled from ≥ 2 experiments, n=10 mice per group, per experiment. * p < 0.05, *** p < 0.001, n.s. not significant.

### IKK2/NFkB signaling interferes with late CD8 T_RM_ transcriptional programming

Recent reports suggest that CD8 T_RM_ development results from a combination of signals that T cells received before tissue entry and signals that occur later at tissue (23, 46-48). In our model, NFkB signals where over-induced after the peak of infection coinciding with a time where some effector antigen specific T cells are being recruited to tissue while others are continuing their differentiation in tissue. We, thus, explored at which time during the contraction phase CD8 T_RM_ loss began in CA-IKK2^ON^ CD8 T cells. For this, we repeated similar experiments to the ones in Fig.1 and monitored the frequency and number of CD69^+^ CA-IKK2^ON^ and control CD8^+^ T_RM_ in lung at day 10 p.i and 30 p.i. (Fig. 4A). We found that the loss of CA-IKK2^ON^ CD8 T_RM_ occcured between day 10 and day 30 p.i. (Fig. 4B, C). The loss of CD8 T_RM_ correlated with a reduction in the number of CD8 T_RM_ expressing the T_RM_ tissue markers CD69 and CD103 (Fig 4C, D). Since we did not observe any defect in the number of circulating memory CD8 T cells in the lung (Fig. 4F) or in the expression of CXCR3, one of the chemokine receptors important for lung recruitment (Fig. 4E)(49), we concluded that NFkB signals must interfere with lung tissue signals that are important for the establishment of CD8 T_RM_.

**Figure 4.**
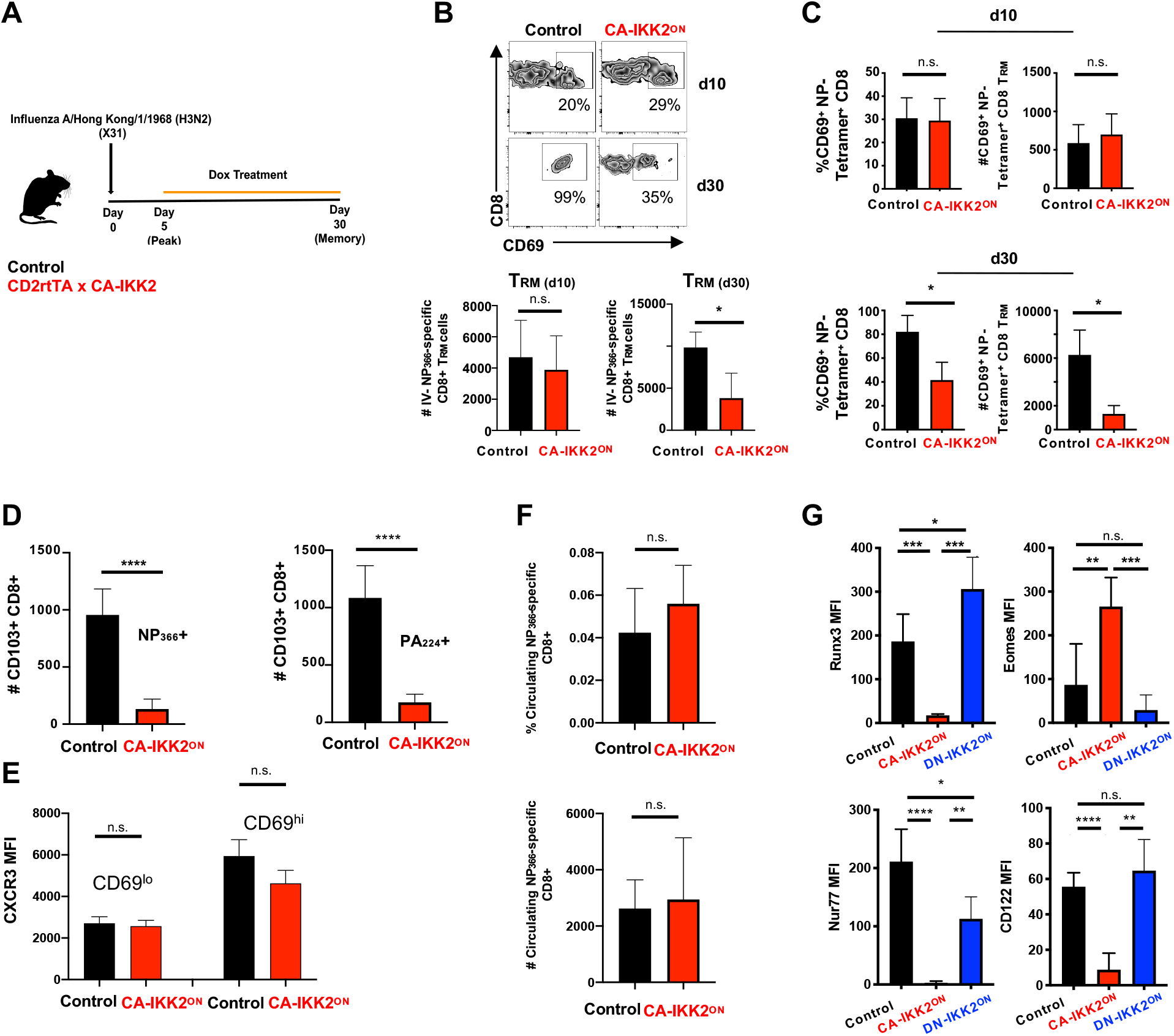
NFkB signaling regulates CD8 T_RM_ transcriptional programming. (A) Experimental Procedure. (B) Groups of control or CD2rtTA x CA-IKK2 mice (n≥3mice per group) were infected with influenza x31 and treated (CA-IKK2^ON^) or not with a doxycycline diet from day 5-30 p.i. Control and CA-IKK2^ON^ Influenza-specific CD8 T_RM_ cells were identified by intravascular labelling and NP_366-374_-specific and PA_224-233_-specific tetramers, and anti-CD8 antibodies by flow cytometry. Dot plot shows frequency of CD8+ CD69+ control and CA-IKK2^ON^ T cells at day 10 and day 30 p.i.. Graph shows number of influenza specific lung T_RM_ cells at day 10 and 30 p.i.. (C) Frequency and number of CD69 positive NP_366-374_-specific CD8 T_RM_ at 10-(top) and 30-days p.i. (bottom) in lung. (D) Number of Influenza specific-CD103+ CD8 T_RM_ cells was determined among NP_366-374_-specific and PA_224-233_-specific CD8+ T cell populations in the lungs 30 days p.i. (E) Expression of CXCR3 in CD69^lo^ and CD69^hi^ populations at day 30 p.i. in lung influenza specific CD8 T_RM_. (F) Frequencies and numbers of circulating, NP_366-374_-specific CD8 T cells determined at day 30 p.i. in lung (G) Expression of lung CD8 T_RM_ – associated transcription factors and CD122 determined at 30 days p.i.Data is representative of ≥ 2 independent experiments. * p < 0.05, ** p < 0.01, *** p < 0.001, **** p < 0.0001, n.s. not significant.

We also assessed whether the defect in the establishment of the CA-IKK2^ON^ CD8 T_RM_ pool was a consequence of T_RM_ transcriptional programming and/or survival. CD8 T_RM_ differentiation requires downregulation of T-box transcription factors Eomes and T-bet (21, 50), induction of Runx3 and Nur77(11) and Blimp-1 in the lung(22) while some studies attribute a role for IL-15 in the homeostasis/survival of CD8 T_RM_ in tissue (26, 51, 52). We observed that CA-IKK2^ON^ CD8 T_RM_ expressed higher levels of T-bet and Eomes than their control counterparts. However, they exhibited reduced levels of Nur77 and Runx3 and normal levels of Blimp-1 (Fig. 4G and *SI Appendix* Fig.S3). Conversely, DN-IKK2^ON^ CD8 T_RM_ exhibited a reversion of the levels of Nur77 and Eomes and an induction of Runx3 over control levels (Fig. 4G). The expression of CD122, one of the chains of the IL-15R, was also impaired in CA-IKK2^ON^ CD8 T_RM_ cells. (Fig.4G). Collectively, these data support the idea that NFkB signaling regulates CD8 T_RM_ transcriptional programming and the imprinting of the CD8 T_RM_ signature.

### NFkB signaling inhibits TGFβ signaling required for CD8 T_RM_ signature molecules

The fact that the number of CA-IKK2^ON^ CD8 T_RM_ cells started to decrease late in the immune response and that this coincided with changes in T_RM_ associated transcription factors (Eomes and Runx3) that are regulated by tissue cues (24, 46), led us to hypothesize that NFkB signaling could be inhibiting tissue signals that are required for lung CD8 T_RM_ differentiation. TGFβ is a crucial cytokine for tissue resident memory differentiation and plays a specific role in lung. Moreover, both Runx3 and CD103 are downstream targets of TGFβ signaling (53, 54), and our data showed defects in both T_RM_ markers in CA-IKK2^ON^ CD8 T_RM_ cells (Fig. 4). Therefore, we tested whether NFkB signaling could inhibit TGFβ signaling in differentiating CD8 T cells (Fig. 5). For this, we used two inducible models where IKK2 signaling can be increased in CD8 T cells. CD8 T cells that were exposed to TGFβ while NFkB signaling was over-induced indeed expressed lower levels of CD103 and Runx3 than TGFβ controls (Fig. 5A). Furthermore, increasing NFkB signaling also resulted in defective canonical (phosphorylated Smad2/3) and non-canonical (phosphorylated ERK) TGFβ signal transduction (Fig 5B,C, *SI Appendix* Data Fig.S4). In other cell types, NFkB can interfere with TGFβ signals through the expression of the inhibitory protein Smad7(55). Thus, we determined the levels of Smad7 under the different conditions. Consistent with the idea that NFkB signals can induce Smad7 in T cells and thereby inhibit TGFβ signaling, we observed that CD8 T cells under high NFkB signaling upregulated Smad7 expression as p-Smad2/3 and pERK levels decreased (Fig. 5C). Collectively, these data shows that IKK2/NFkB signaling can negatively regulate TGFβ signaling and the CD8 T_RM_ markers, Runx3 and CD103.

**Figure 5.**
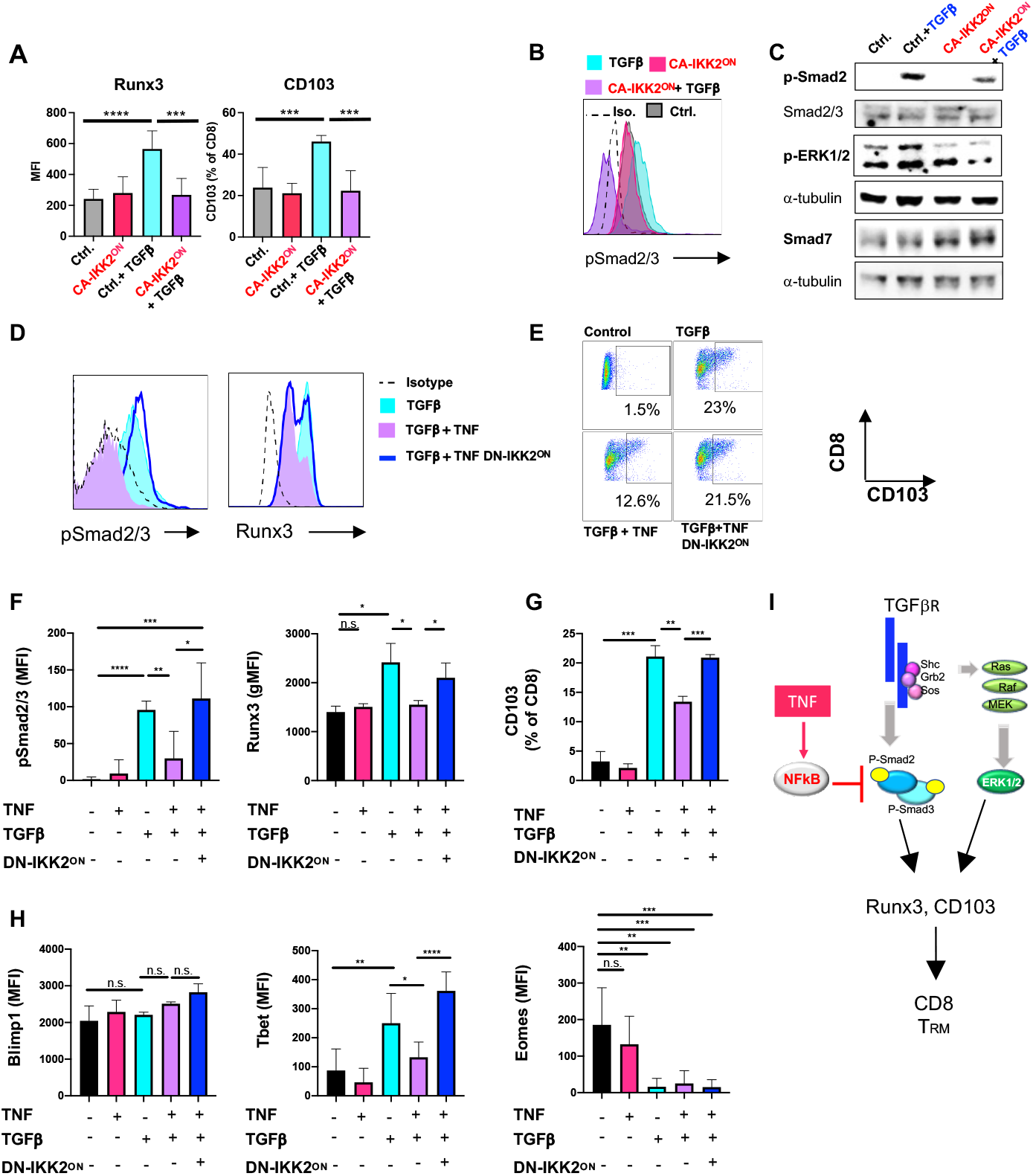
TNF and NFkB signaling inhibit TGFb signaling and downstream T_RM_ markers, CD103 and Runx3. (A-C) Splenocytes from OT-1 (control) or OT-1xIKK2^fl/fl^xGzB^Cre^ mice (CA-IKK2^ON^ samples) were stimulated for 48 hours with OVA peptide to generate effector CD8 T cells that were next stimulated with TGFβ (A) Expression of Runx3 and CD103 (MFI) were determined 24 hours after. (B) Histograms show phosphorylated levels of Smad2/3 30 minutes following TGFβ stimulation on control (blue) or CA-IKK2^ON^ CD8 T cells. (C) phospho-Smad2 and -ERK1/2, expression of Smad7 and loading control tubulin were determined by immunoblot with specific antibodies after 30 minutes of TGFβ stimulation. (D-I) Splenocytes from CD2rtTA x DN-IKK2 and control mice were stimulated with anti-CD3/ CD28-specific antibodies. 1 day later, cells were treated or not with TNF and doxycycline for another 24 hours. TGFβ was then added to indicated samples. Histograms show phospho-Smad2/3 and Runx3 levels on CD8 T cells 30 minutes post-TGFβ addition (D). (E) Dot plots show frequency of CD8+ CD103+ CD8 T cells in the conditions shown. (F-H) Levels of phospho-Smad2/3, Runx3, Blimp-1, Tbet and Eomes as well as % of CD103^+^ CD8 T cells were determined 24 hours post-TGFβ stimulation. (i) Working model. Data shown is pooled from ≥ 2 experiments of n≥ 3 independent experiments. * p < 0.05, ** p < 0.01, *** p < 0.001, **** p < 0.0001, n.s. not significant.

### TNF-mediated NFkB signaling inhibits TGFβ signaling for T_RM_

NFkB signaling is a mediator of inflammatory signals during respiratory infections(56-58). One of the most understood NFkB triggers is the pro-inflammatory cytokine TNF, which has been associated to T_RM_ in the lung (59) and signals through the canonical NFkB pathway(35). Based on these, we hypothesized that TNF could, via NFkB, inhibit TGFβ signaling and impair CD8 T_RM_. To assess this, we designed an experiment where CD8 T cells differentiating in the presence of TNF were exposed to TGFβ and then, measured changes in TGFβ signaling as well as T_RM_ markers downstream of TGFβ, Runx3 and CD103. As expected, TNF did not induce TGFβ signaling or the expression of Runx3 and CD103 while TGFβ did. However, consistent with the idea that TNF inhibits TGFβ signaling, CD8 T cells differentiating in the presence of TGFβ were impaired in the phosphorylation of Smad2/3 and the expression of Runx3 or CD103 when TNF was present (Fig. 5D-G).

To confirm whether the ability of TNF to inhibit TGFβ signaling was NFkB dependent, we repeated the same experiments with DN-IKK2^ON^ CD8 T cells and use doxycycline to inhibit NFkB signaling. Confirming our hypothesis, DN-IKK2^ON^ CD8 T cells remain unresponsive to the effects of TNF on TGFβ signaling (Fig. 5D-G). Thus, TNF inhibits TGFβ signaling and proteins crucial for CD8 T_RM_ via NFkB.

We also assessed whether TNF and NFkB signaling could affect the induction of other transcription factors important for lung CD8 T_RM_. Blimp-1 was not affected by TNF and/or TGFβ. By contrast, T-bet and Eomes were. Remarkably, Eomes expression was inhibited by TGFβ (60) and neither TNF nor inhibiting NFkB signaling could revert it (Fig. 5H). In summary, these results support the idea that pro-inflammatory cytokines able to induce NFkB, such as TNF, can inhibit TGFβ signals in CD8 T cells and, thereby, impair their differentiation towards the CD8 T_RM_ fate (Fig. 5I).

### At memory, NFkB signaling promotes lung CD8 T_RM_ survival

Upon influenza infection, lung CD8 T_RM_ cells fail to persist weakening protective immunity against the same or other influenza variants (18, 61). In prior work using NFkB pharmaceutical inhibitors, it was found that once memory CD8 T cells are generated their maintenance depended on NFkB signals(42). Although these studies did not distinguish between the different T cell memory subsets, the data conflicted with our findings that enhanced NFkB signaling is detrimental to CD8 T_RM_ differentiation (Fig. 1 and 4). Therefore, we decided to address the impact of NFkB signaling on CD8 T_RM_ at memory once memory T cells have been formed. For this, we used the CA-IKK2^ON^ inducible model and allowed the generation of CD8 T_RM_ in the lung upon IAV infection. 30 d.p.i, we compared control and doxycycline-treated mice for changes in the number of CD8 T_RM_ in the lung. We observed a ∼ 6-fold increase in the number of IAV specific CD8 T_RM_ cells in the lung when NFkB signaling had been induced (Fig. 6A). Strikingly, this was the opposite effect that increasing NFkB signals during contraction had in CD8 T_RM_ generation (Fig. 1). The increase in CA-IKK2^ON^ CD8 T_RM_ at memory correlated with higher levels of CD122 and Bcl-2, suggesting NFkB signals at memory mediate CD8 T_RM_ survival (Fig. 6B).

**Figure 6.**
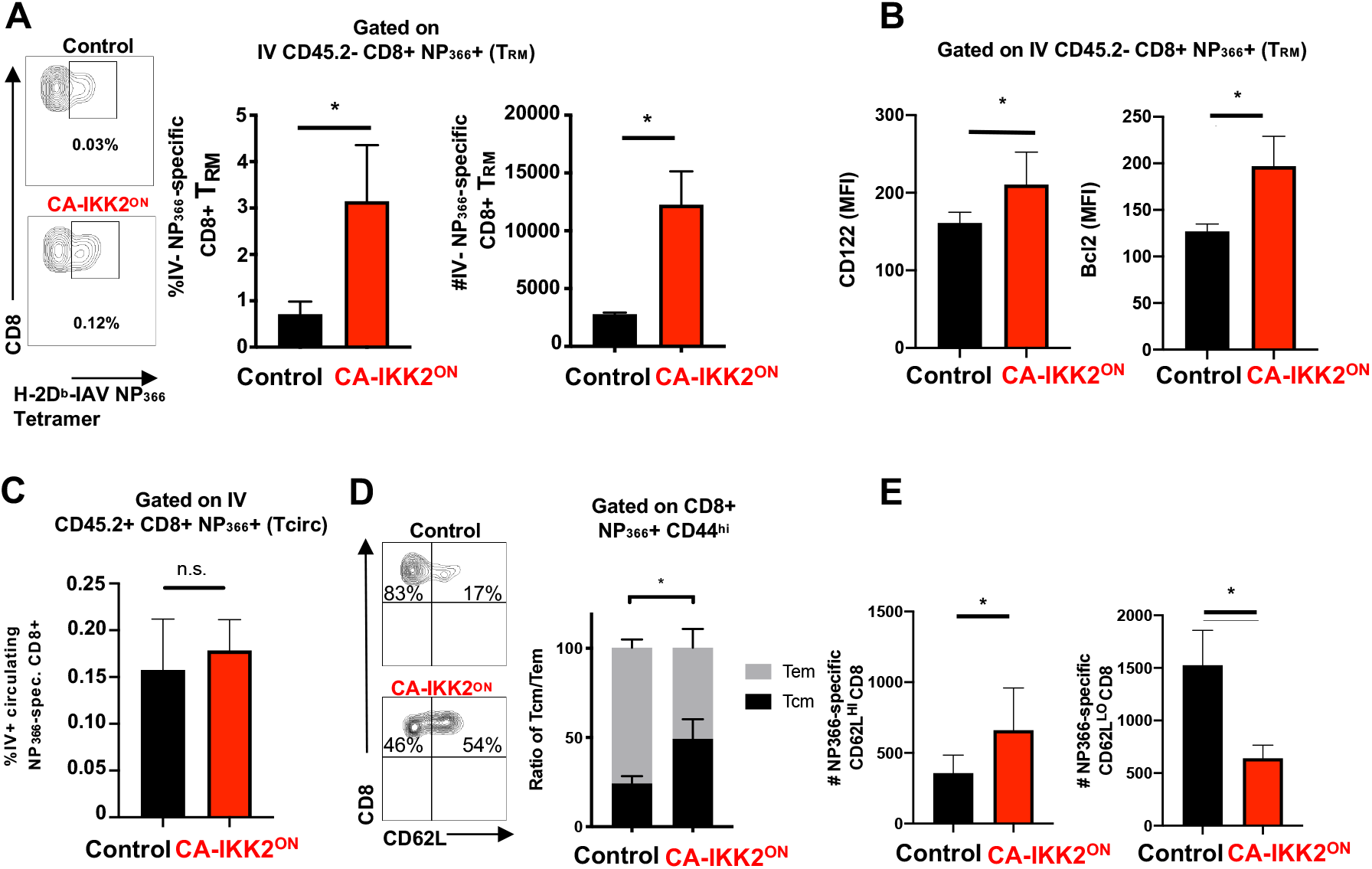
Increasing NFkB signaling at memory improves CD8 T_RM_ survival. Groups of control or CD2rtTA x CA-IKK2 mice (n ≥ 3 mice per group) were infected with influenza x31. At day 30 p.i. mice were fed a doxycycline (CA-IKK2^ON^) or a control containing diet for 15 days. (A) Frequencies and numbers of influenza specific CD8 T_RM_ identified by intravascular staining in lungs. (B) Bcl2 and CD122 expression on lung CD8 T_RM._ (C) Frequency of circulating NP_366-374_-specific CD8 T cells determined in the lungs by intravascular labeling. (D-E) Frequency, ratio and number of NP_366-374_-specific CD8^+^ T_CM_ (CD8+ Db-NP-tet+ CD44^hi^ CD62L^hi^) and T_EM_ in mediastinal lymph nodes. (Representative data shown from ≥ 2 experiments. * p < 0.05, n.s. not significant.

In most tissues T_RM_ maintenance is independent of the input of circulating memory T cells (62, 63) although, in the lung this is still controversial (61, 64). Curiously in our studies, increasing NFkB signals at memory only boosted the frequency of IAV specific CD8 T_RM_ in the lung. The frequency of IAV specific circulating memory T cells remain unaltered (Fig. 6C), suggesting NFkB signals improve CD8 T_RM_ maintenance in the lung by supporting CD8 T_RM_ survival.

Finally, NFkB signaling at memory was also beneficial for the CD8 central memory pool as we also observed a significant increase in the number of IAV specific CD8 T_CM_ in draining lymph nodes after doxyclycline treatment (Fig. 6. D, E). Together, these data reveal that NFkB signaling differentially affects tissue resident memory depending on the stage of differentiation of the CD8 T cell (before and after becoming CD8 T_RM_). Most importantly, our results support the idea that enhancing NFkB signaling in CD8 T cells once the T_RM_ pool has been established could improve CD8 T_RM_ maintenance in tissue.

## Discussion

T cell memory in tissues is an essential part of mucosal immunity that protects against infection and disease. Here we show that the pro-inflammatory signaling pathway IKK2/NFkB, is critical for both the generation and maintenance of the CD8 T_RM_ pool after infection. Our results, mainly refer to influenza specific CD8 T_RM_ in the lung, a tissue where maintaining a long-lived CD8 T_RM_ pool is crucial for protective immunity(18) but challenging, due to the CD8 T_RM_ short life-span (65). The reasons why resident memory CD8 T cells are short-lived in the lung but not in other tissues are still unclear. However, our work suggests that boosting NFkB signaling at the end of the immune response might offer a therapeutic avenue to increase CD8 T_RM_ survival and protective immunity upon infection or vaccination. Furthermore, since improved survival also occurred for CD8 T_CM_ of the draining lymph nodes, this suggests a controlled increase in NFkB signaling could boost several subsets of the T cell memory pool.

It is striking that the same signaling pathway, NFkB, operates in opposite manners for CD8 T cells depending on their differentiation stage (during T_RM_ differentiation and at memory). This could be due to epigenetic modifications that regulate the accessibility of NFkB to specific gene loci depending on the differentiation stage of a CD8 T cell. Alternatively, changes in the environmental cues as the infection resolves could also explain the differential impact on CD8 T_RM_ when levels of NFkB signaling increase. Although our findings cannot distinguish between these two possibilities, our data suggest that NFkB signaling does interfere with TGFβ to skew CD8 T effectors away from T_RM_. TGFβ is a universal driver of CD8 T_RM_ whose levels in tissue can change depending on age and in the context of diseases such as infection, autoimmunity, asthma, or fibrosis(66). Importantly, multiple signals can also locally trigger the induction of NFkB signaling in CD8 T cells, including antigen, TLRs and pro-inflammatory cytokines(35-37). We show that TNF, a known driver of NFkB (35) is able to inhibit TGFβ b-dependent signaling and T_RM_ programming in CD8 T cells. TNF and other pro-inflammatory cytokines such as IL-6 are heavily produced in pathological settings of chronic inflammation and could easily affect the levels of NFkB signals(67-69) a CD8 T cell experience in tissue. For the CD8 T cells that also encounter TGFβ locally, this could decrease their likelihood to become CD8 T_RM_. Further evaluation of the dynamics of CD8 T_RM_ during infections with a strong pro-inflammatory profile could provide insight into how overt inflammation may affect long-term immunity and inform of specific therapeutics targeting NFkB or its pro-inflammatory drivers to either boost or deplete tissue resident memory. This might be especially relevant in the context of immune treatments that are linked to high levels of inflammation and that in the case of cancer(70, 71) or autoimmunity (RA) have a T_RM_ component that affects disease outcome (14, 72). In the same line, it would also be important to evaluate how current treatments targeting TNF or NFkB signaling in the Clinic, affect the establishment of tissue resident memory in patients.

We also found that NFkB signaling did not affect CD8 T_RM_ in the same manner as it did CD8 T_CM_ or T_EM_ development, indicating that NFkB signaling is a key regulator of T cell memory diversity. Modulation of NFkB signaling levels may serve as an opportunity to regulate specific T cell memory subsets depending on their role in disease. Finally, our findings also underscore the impact that fluctuating levels of a single signaling pathway can have on the quality of T cell memory depending how and when during the infection these levels change. This may be particularly important for pleiotropic signaling pathways such as NFkB where multiple stimuli feed in (including patient’s treatments) and can easily add up to shape T cell fate. Incorporating this concept into current vaccine strategies could aid to improve their long-term efficacy.

## Materials and Methods

### Mice

OT-1Thy1.1+ TCR transgenic strain, C57BL/6J (B6), B6.SJL-*Ptprc*^*a*^ *Pepc*^*b*^/BoyJ (CD45.1 congenic C57BL/6), B6.Cg-*Gt(ROSA)26Sor*^*tm4(Ikbkb)Rsky*^/J (IKK2-CA^fl/fl^), B6 -Tg (GzB-cre)1Jcb/J (GzB-Cre) mice (Jackson Laboratory, Bar Harbor, ME) along with OT-1Thy1.1XIKK2CA^fl/fl^ xGzB^Cre^; CD2rtTA x CA-IKK2 (tetracycline-inducible constitutive active IKK2) and CD2rtTA x DN-IKK2 (tetracycline-inducible dominant negative IKK2) mice were maintained under specific pathogen-free conditions at the University of Missouri. All mouse strains were screened for transgene homozigocity by PCR. Mice were aged between 8-13 weeks at the time of infectin. Infection and maintenance of mice infected with influenza virus or vesicular stomatitis virus occurred in an ABSL2 facility at the University of Missouri. All animal procedures were conducted according to the NIH guidelines for the care and use of laboratory animals and were approved by the University of Missouri Institutional Animal Care and use Committee.

### Reagents and antibodies

Biotinylated, influenza-specific monomers (H-2Db NP366-374 ASNENMETM, H-2D^b^ PA224-233 SSLENFRAYV, I-A^b^ NP311-325 QVYSLIRPNENPAHK) were obtained from the NIH Tetramer Core Facility (Atlanta, GA). Biotinylated H-2K^b^ monomers for OVA (SIINFEKL) and VSV NP52-59 (RGYVYQGL) were generated in our laboratory. Biotinylated monomers were tetramerized using fluorescently labeled streptavidin (Biolegend, San Diego, CA). Doxycycline containing diet (6 g/kg) was purchased from Envigo (Indianapolis, IN).

For Flow cytometry we used antibodies anti-CD8 (53-6.7), CD4 (L3T4), CD45.2 (104), CD44 (IM7), CD62L(MEL-14), TNF (MP6-XT22) and CXCR3 (CXC3-173) from Biolegend; anti-CD103 (M290), CD69 (H1.2F3), CD122 (TM-b1), Blimp1 (6D3),T-bet (O4-46), IFN-g(XMG1.2) and Phospho-SMAD2/3 from BD Biosciences; anti-Runx3 (527327) from R&D systems; anti-Eomes (Dan11mag) from Thermo Scientific and anti-Luciferase from Rockland, Inc. For western blotting and immunostaining we used antibodies anti-phosphorylated-SMAD2(Ser465/Ser467) (E8F3R), SMAD2/3 (D7G7), phosphorylated ERK1/2 from Cell Signaling; anti-SMAD7(293739) from R&D Systems; anti-α-tubulin (B-5) from Sigma and secondary antibodies goat anti-mouse and anti-rabbit from Li-Cor Biotechnology.

### Virus infections

Mice were infected intranasally with 1000 pfu Influenza A/HKx-31 (X31,H3N2) for sublethal infection or intravenously with vesicular stomatitis virus (VSV) (2×10^6^ pfu), unless otherwise indicated in Figure legends. For heterologous infection experiments, mice were primed intranasally with 5×10^4^ pfu VSV-OVA, then challenged 30 days later with 5000 pfu of influenza A/PR8-OVA (PR8, H1N1).

### Viral titers

The TCID_50_ of influenza virus was determined using MDCK cells as described (*74*). Briefly, lung samples were homogenized using a Mini-BeadBeater (BioSpec, Bartlesville, OK) and cleared homogenate was used to inoculate confluent MDCK cell monolayers. 24 hours post inoculation, the supernatant was discarded and replaced with fresh media (DMEM containing 0.0002% Trypsin). Agglutination of chicken RBCs (Rockland Immunochemicals Inc., Limerick, PA) was utilized to determine the presence of influenza virus after 3 days of culture.

### *In vivo* antibody labeling and flow cytometry

For *in vivo* antibody labeling and differentiation of T cells circulating in the vasculature or resident in parenchyma (TRM) tissues, three minutes before being killed, mice were injected intravenously via tail vein injection with 2 mg PE-labeled CD45.2 (clone 104) or PE-labeled CD8b, (clone Ly-3). Lungs, kidney, spleen, and mediastinal lymph node tissues were harvested, and lymphocytes isolated. Next, lymphocytes were stained *in vitro* with anti-CD8α antibodies along with antibodies to other surface and intracellular markers conjugated to fluorochromes. Stained cells were run on a LSR Fortessa flow cytometer (BD, San Jose, CA), and analyzed using with FlowJo software (Tree Star, Inc., Ashland, OR).

### Intracellular cytokine staining

Lymphocytes were isolated from the lungs of VSV-OVA-challenged mice and stimulated *ex vivo* with OVA peptide 1 mM) in the presence of Golgi-Plug (BD Biosciences) for 5 hours. Following incubation, cells were harvested and antigen specific CD8^+^ T cells were assessed for the expression of TNF and IFNg by flow cytometry.

### CD8 T cell enrichment and adoptive transfer

Splenocytes were harvested from CD2rtTA x CA-IKK2 mice and polyclonal CD8 T cells were purified by magnetic selection (CD8 T cell isolation kit by Miltenyi Biotech Auburn, CA). 5×10^5^ CD8 polyclonal or 10^4^ OT-1Thy1.1XIKK2CA^fl/fl^ xGzB^Cre^ monoclonal naïve T cells were adoptively transferred into congenic C57BL/6 mice 1 day prior to intranasal infection with influenza or VSV-OVA virus respectively.

### *In vitro* culture and cytokine stimulation

Splenocytes isolated from OT-1Thy1.1XIKK2CA^fl/fl^ xGzB^Cre^ mice were stimulated with 20 nM OVA peptide for 48 hours at a concentration of 1×10^6^ cells/ml. TGFβ (R&D Systems, Minneapolis, MN) was then added to a final concentration of 50 ng/ml. At 30 minutes and 24 hours post TGFβ stimulation, cells were harvested for analysis by flow cytometry and western blotting. Splenocytes from CD2rtTA x DN-IKK2 mice were stimulated *in vitro* at 1×10^6^ cells/ml with 10 mg/ml anti-CD3 (clone 145-2C11) and 10 mg/ml anti-CD28 (clone 37.51) (ThermoFisher Scientific, Waltham, MA). Following 24 hours of stimulation, cells were divided and incubated in the presence or absence of 125 ng/ml recombinant TNF (R&D Systems, Minneapolis, MN) for 24 additional hours. The cells were again divided and incubated in the presence or absence of 50 ng/ml TGFβ (R&D Systems, Minneapolis, MN). Cells were harvested at 30 minutes and 24 hours post addition of TGFβ and analyzed by flow cytometry.

### Western blotting

*In vitro* stimulated cells were lysed in lysis buffer containing 10mM HEPES, 10mM KCl, 0.1mM EDTA, 0.2mM EGTA, 0.5% NP40, 1mM DTT, 2 mM Na_3_VO_4_, 20 mM NaF, 1 mg/ml Leupeptin, 1 mg/ml Aprotonin, 1 mM PMSF. Samples were resolved on a 10% SDS-PAGE gel and transferred to nitrocellulose membrane. Membranes were blocked with Blotting Grade Blocker (Bio-Rad, Hercules, CA) and probed with specific primary and secondary antibodies. Blots were imaged on a Li-Cor Odyssey XF (Li-Cor, Lincoln, NE) and analyzed using Image Studio (Li-Cor, Lincoln, NE).

### Statistical analysis

Statistical analysis was performed using the Prism software (GraphPad). Data are presented as mean +/-standard deviation. Statistical significance to compare two quantitative groups was evaluated using a two-tailed t test. A value for significance of p< 0.05 was used throughout the study, and statistical thresholds of 0.05, 0.005, 0.0005, as well as 0.0001 are indicated in the figures signified via asterisks (as described in the figure legends).

## Acknowledgments

We thank Dr. Nick Goplen and Dr. Vikas Saxena for discussion and Bianca Gordon for screening. We thank Dr. Zamoyska, Dr. Bruce Richardson, and Faith Strickland for generously providing us with CD2rtTA mice. Bernd Baumann, Dr. Thomas Wirth for generously providing us with IKK2-CA and IKK2-DN mouse strains. MU Office of Research and School of Medicine animal vivarium staff for assistance with mice. Kathy Schreiber and Daniel Jackson for assistance in the Flow Cytometry Core Facility. Supported by National Institutes of Health grants R01 AI110420-01A1 and R56 AI110420-06A1, 1U01CA244314 as well as Internal funding from the School of Medicine (ET).

## Notes

### Competing Interest Statement

The authors have declared no competing interest.

