## Supplemental material Fig S1-S4 for "IKK2/NFkB signaling controls lung resident CD8 T cell memory during influenza infection"

**Supplemental information Appendix**

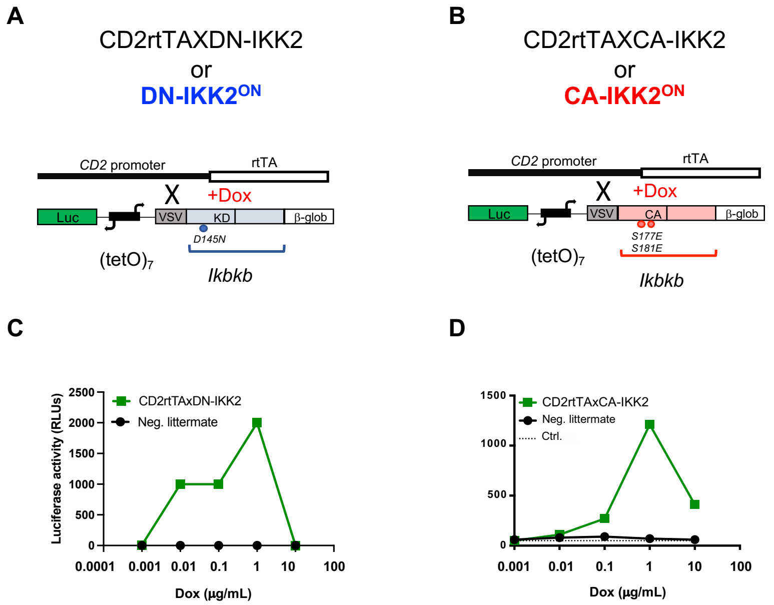


**Fig.S1 related to Fig.1. T cell restricted Tet ON inducible models to inhibit or enhanced IKK2 enzymatic activity** (**A,B**) Genetic strategy to generate T cell restricted inducible models DN-IKK2$^{ON}$ (**A**) and CA-IKK2$^{ON}$ (**B**) from crossing CD2rtTA mice generated in (41) and CA-IKK2 and DN-IKK2 described in (39, 40). The tetracycline-responsive transactivator domain rtTA is constitutively expressed as a transgene in the T cell lineage driven by human CD2 regulatory regions. The rtTA-activated promoter (tetO)7 directs the transcription of luciferase and Ikbkb that has been mutated in the kinase domain (KD, D145N; CA, S177E, S181E). VSV, vesicular stomatitis tag; b-glob, b-globin intron/poly(A) signal. In mice resulting from the cross of CD2rtTA and CA-IKK2 or DN-IKK2 strains constitutive and death kinase forms of the kinase IKK2 are expressed only in the presence of tetracycline or its derivatives (doxycycline) in cells of the T cell lineage. (**C, D**) CD2rtTAxDN-IKK2 or CD2rtTAxCAIKK2 splenocytes or their negative littermates were stimulated with anti-CD3/28 and different doses of doxycycline (dox). Luciferase activity was measured on day 2.

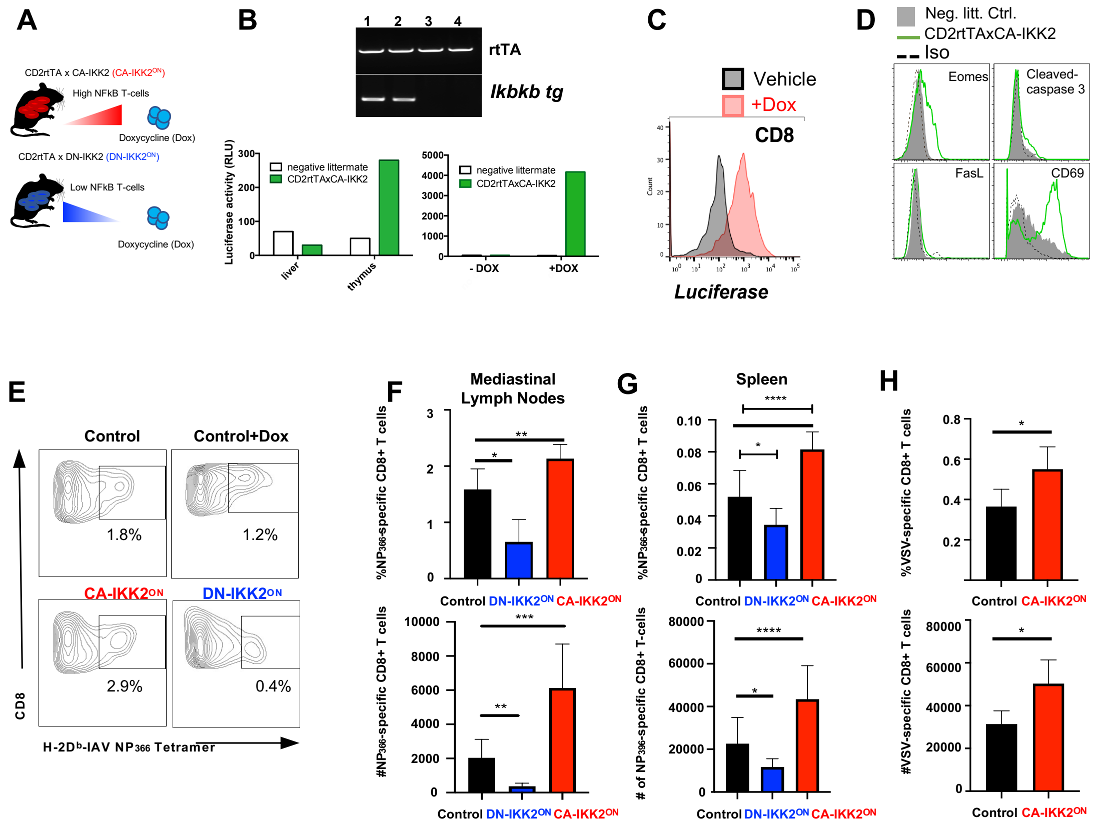


**Fig.S2 related to Fig.1. Enhancing NFkB signals at the end of the immune response boosts the generation of memory CD8 T cells. (A).** TetON Mouse inducible models to enforce (CD2rtTA x CA-IKK2) or inhibit (CD2rtTA x DN-IKK2) IKK2/NFkB signaling. (**B**) PCR based analysis of indicated genes from CD2rtTA x CA-IKK2 mouse tails (lanes 1,2) and parental negative controls (lanes 3,4). Left graph: liver and thymus from CD2rtTA x CA-IKK2 mice treated with 4 mg/mL doxycycline solution were lysed, and luciferase activity measured to indicate absence of expression of the transgene in non-lymphoid tissues (liver) versus lymphoid tissue (thymus). Right graph: Purified T cells from CD2rtTA x CA-IKK2 mice were stimulated with anti-CD3/CD28 antibodies in varying concentrations of dox. Luciferase activity was assessed in an enzymatic assay and indicates dose-dependent induction of the transgene. (**C**) IKK2 constitutive activation correlates with induction of Luciferase-reporter expression measured by flow with a Luciferase-specific antibody in CD8 T cells from the lymph nodes of CA-IKK2$^{ON}$ mice that have been treated with doxycycline containing chow for 25 days. (**D**) Expression of markers indicated in CD8 T cells of CD2rtTA x CAIKK2 mice (green) or negative littermates (grey) that have been treated with dox solution for 7 days. (**E-H**). Groups of control, CD2rtTA x CA-IKK2 (CA-IKK2$^{ON}$), or CD2rtTA x DN-IKK2 (DN-IKK2$^{ON}$) mice (n=3 mice per group) were infected with the x31 strain of influenza virus (IAV-x31). Beginning at day 5 post-infection (p.i.), negative littermates (control + dox) and inducible mice were fed a doxycycline containing diet or control diet (control). NP366-374-specific CD8 T cells (Db-NP-tet+, CD8+ CD44$^{hi}$) were identified in the mediastinal lymph nodes after day 30 p.i. (**E, F**) and the spleen (**G**) by flow cytometry. (**H**) Groups of control or CD2rtTA x CA-IKK2 (CA-IKK2$^{ON}$) mice were infected with VSV. Mice were fed a doxycycline-containing diet from days 5 – 30 p.i. and VSV-specific memory CD8 T cells (Kb-N-tet+ CD8+ CD44$^{hi)}$ were assessed in lymph nodes by flow cytometry. Representative data shown from among 3 individual experiments. * $p < 0.05$, ** $p < 0.01$, *** $p < 0.001$, **** $p < 0.0001$.

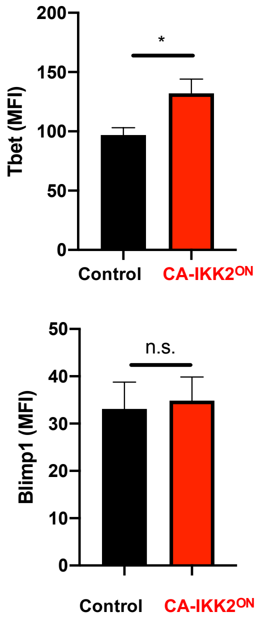


**Fig. S3 related to Fig.4. NF-kB-mediated regulation of tissue resident memory-associated transcription factors.** Groups of control or CD2rtTA x CA-IKK2 (CA-IKK2$^{ON}$) mice (n≥3mice per group) were infected with the X31 strain of influenza virus. Beginning at day 5 p.i., mice were fed a doxycycline containing diet or control diet. Influenza-specific CD8 $T_{RM}$ were identified by intravascular staining with PE-labeled CD45.2 antibody. Expression of Tbet and Blimp1 was determined *ex vivo* after 30 days p.i. by flow cytometry. Data is representative of ≥ 2 independent.

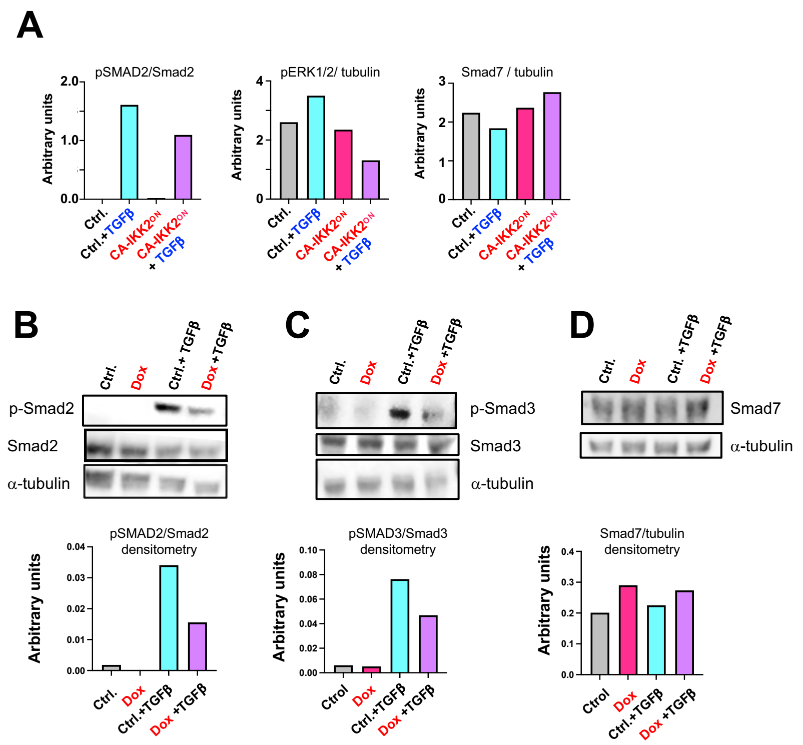

**Fig. S4 related to Fig.5. NFkB inhibits TGFb signaling in the tetON CAIKK2$^{ON}$ inducible model. (A)** Graphs show densitometry of phospho-Smad2, phospho-ERK1/2 and Smad7 for immunoblots shown in Fig. 5C. Representative of 2 independent experiments **(B-D)** Immunoblot and densitometry of phospho-Smad2, -Smad3 and Smad7 using T cells from the CD2rtTAxCA-IKK2 tetON inducible model. Dox indicates samples where cells were treated with doxycycline to induce constitutive active IKK2 activity. Cells were CD2rtTAxCA-IKK2 CD8 T cells were stimulated as in Fig.5C in the presence or absence of TGFβ. In brief, splenocytes from CD2rtTAxCA-IKK2 mice were stimulated for 24 hours with antibodies anti-CD3 and -CD28 (10 mg/ml). Dox (1 mg/mL) was added or not (control) for an additional 24 hours. TGFβ was added to a final concentration of 50 ng/ml. Data shown is representative experiment of 3 independent experiments.
